# Deep proteomics network and machine learning analysis of human cerebrospinal fluid in Japanese encephalitis virus infection

**DOI:** 10.1101/2022.06.19.496758

**Authors:** Tehmina Bharucha, Bevin Gangadharan, Abhinav Kumar, Ashleigh C. Myall, Nazli Ayhan, Boris Pastorino, Anisone Chanthongthip, Manivanh Vongsouvath, Mayfong Mayxay, Onanong Sengvilaipaseuth, Ooyanong Phonemixay, Sayaphet Rattanavong, Darragh P. O’Brien, Iolanda Vendrell, Roman Fischer, Benedikt Kessler, Lance Turtle, SEAe collaborators, Xavier de Lamballerie, Audrey Dubot-Peres, Paul N. Newton, Nicole Zitzmann

## Abstract

Japanese encephalitis virus (JEV) is a mosquito-borne flavivirus, and leading cause of neurological infection in Asia and the Pacific, with recent emergence in multiple territories in Australia in 2022. Patients may experience devastating socioeconomic consequences; JEV infection (JE) predominantly affects children in poor rural areas, has a 20-30% case fatality rate, and 30-50% of survivors suffer long-term disability. JEV RNA is rarely detected in patient samples, and the standard diagnostic test is an anti-JEV IgM ELISA with sub-optimal specificity; there is no means of detection in more remote areas. We aimed to test the hypothesis that there is a diagnostic protein signature of JE in human cerebrospinal fluid (CSF), and contribute to understanding of the host response and predictors of outcome during infection.

We retrospectively tested a cohort of 163 patients recruited as part of the Laos central nervous system infection study. Application of liquid chromatography and tandem mass spectrometry (LC-MS/MS), using extensive offline fractionation and tandem mass tag labelling, enabled a comparison of the CSF proteome in 68 JE patient vs 95 non-JE neurological infections. 5,070 proteins were identified, including 4,805 human proteins and 265 pathogen proteins. We incorporated univariate analysis of differential protein expression, network analysis and machine learning techniques to build a ten-protein diagnostic signature of JE with >99% diagnostic accuracy. Pathways related to JE infection included neuronal damage, anti-apoptosis, heat shock and unfolded protein responses, cell adhesion, macrophage and dendritic cell activation as well as a reduced acute inflammatory response, hepatotoxicity, activation of coagulation, extracellular matrix and actin regulation. We verified the results by performing DIA LC-MS/MS in 16 (10%) of the samples, demonstrating 87% accuracy using the same model. Ultimately, antibody-based validation will be required, in a larger group of patients, in different locations and in field settings, to refine the list to 2-3 proteins that could be harnessed in a rapid diagnostic test.

**Author summary:** Japanese encephalitis virus (JEV) is a leading cause of brain infection in Asia and the Pacific, with recent introduction in multiple territories in Australia in 2022. Patients may experience devastating socioeconomic consequences; JEV infection (JE) predominantly affects children in poor rural areas, has a 20-30% case fatality rate, and 30-50% of survivors suffer long-term disability. The disease is difficult to diagnose, and there are no rapid tests that may be performed in remote areas that it exists such that we remain unclear of the burden of disease and the effects of control measures. We aimed to apply a relatively novel method to analyse the proteins in patients with JE as compared to other neurological infections, to see if this could be useful for making a diagnosis.

We tested the brain fluid of 163 patients recruited as part of the Laos central nervous system infection study. We used a method, ‘liquid chromatography mass spectrometry’ that does not require prior knowledge of the proteins present, that is you do not target any specific protein. Over 5,000 proteins were identified, and these were analysed by various methods. We grouped the proteins into different clusters that provided insight into their function. We also filtered the list to 10 proteins that predicted JE as compared to other brain infections. Future work will require confirmation of the findings in a larger group of patients, in different locations and in field settings, to refine the list to 2-3 proteins that could be harnessed in a rapid diagnostic test.

## Introduction

Japanese encephalitis virus (JEV) is a mosquito-borne flavivirus, and a leading cause of neurological infection as Japanese encephalitis (JE) in Asia. It is of considerable public health importance, with recent estimates based on sparse data suggesting 1.5 billion people at risk with 42,000 cases per year (1, 2). It is an emerging disease, with recent evidence of JEV in multiple territories in Australia (3). Patients may experience devastating socioeconomic consequences; JE predominantly affects children in poor rural areas, has a 20-30% case fatality rate, and 30-50% of survivors suffer long-term disability (4). Although no specific treatment is available, several vaccines are available and recommended by the WHO (5, 6). Although recent efforts have strengthened JEV vaccination programs, still only 15 of 24 endemic countries include JEV vaccine in routine immunisation policies, and even then, it is not uniformly nationwide, with vaccine coverage in targeted areas reported to be as low as 39% (7). JEV is a zoonosis, and sustained vaccine coverage is essential to control disease.

A fundamental limitation in the control of JE is the poor accuracy of existing diagnostic tests, requirement for lumbar puncture and laboratory capacity for diagnosis (8). Surveillance data suggest that only 11 of 24 countries meet the minimum surveillance standards, equivalent to diagnostic testing in a sentinel site (7). This is a threat to vaccine implementation, as accessible and accurate diagnostics are essential to understand epidemiology, effectiveness of vaccination, identify associated research knowledge gaps and facilitate public engagement. This also has implications for appropriate risk-assessment for travellers. Aside from JEV control, diagnosis is crucial for patients, families and health-workers, to be able to institute appropriate supportive and rehabilitation care, stop unnecessary antibiotics, or if the test is negative to prompt further investigation.

The gold-standard JEV test is a neutralisation assay. However this requires paired acute and convalescent sera, is laborious, time-consuming, requires specialist skills, high-level isolation facilities for viral cell culture and may not define the infecting virus in secondary flavivirus infections (8). The WHO recommended diagnostic test is anti-JEV IgM antibody capture ELISA (MAC-ELISA) of cerebrospinal fluid (CSF). There are limited data from field studies comparing CSF MAC ELISA with neutralisation assays. The manufacturer of the only available commercial kit for clinical diagnosis (InBios) quotes a sensitivity of >90% for well-characterised CSF samples, but sensitivity in the field is as low as 53% (9). There are also increasingly recognised problems with specificity related to prior vaccination and cross-reactivity with other flaviviruses (10, 11). Reported specificity is >90%, however a study by our group demonstrated that 13% of patients with JE IgM detected in CSF by MAC-ELISA had another pathogen detected that may have explained the presentation (10).

Detection of JEV RNA would be highly specific, but the period of viraemia is brief and hard to capture clinically, often occurring before the onset of neurological symptoms and signs. RT-qPCR remains insensitive irrespective of the analytical sensitivity or gene targets (12). For this reason, the application of metagenomics is not likely to significantly improve JEV RNA detection.

Uniquely untargeted and powerful, the application of liquid-chromatography mass-spectrometry (LC-MS) proteomics techniques to clinical samples represent a relatively novel approach to improve diagnosis of JE (13, 14). Such an approach is based on the hypothesis that there is a protein signature in CSF specific for JE, and that this could be harnessed in an antibody-based point-of-care test. Furthermore, deep proteomics exploration provides insights into disease processes and potential therapeutic targets. Network science and machine learning are two complementary disciplines enabling insights into complex high dimensional data (15, 16). Networks, comprised of nodes and links, are naturally attuned to problems where features have a relational structure (17) and have a track record of success in understanding networks of biological interactions (18). On the other hand, machine learning can uncover signals in data related to outcome variables and identify predictive markers of disease, a vital exploratory process for constructing diagnostics (19). Used in conjunction, network science and machine learning provide novel characterisation of disease states and can identify robust predictive markers of disease (20).

Herein we aimed to test the hypothesis that there is a diagnostic protein signature of JE by performing LC-MS/MS in patient samples recruited as part of the Laos CNS study, incorporating differential expression, network and machine learning analysis. A subsidiary aim was to utilise the data in the same workflow to evaluate proteins associated with outcome of JE. We first performed a pilot feasibility study (n=15) and then in a larger verification study (n=148) including a sample size based on a power calculation. These data were combined in the final analysis. The results were verified by performing data independent acquisition (DIA) LC-MS/MS in 16 (10%) of the samples.

## Materials and methods

### Patient samples

A prospective study of central nervous system (CNS) infection has been conducted at Mahosot Hospital, Vientiane, Laos, since 2003. Methods and results from 2003-2011 have been described (21). Patients from 2014-2017 were part of the Southeast Asia Encephalitis Project (22). Inpatients of all ages were recruited for whom diagnostic lumbar puncture was indicated for suspicion of CNS infection because of altered consciousness or neurologic findings and for whom lumbar puncture was not contraindicated. There was no formal definition for CNS infection; patient recruitment was at the discretion of the responsible physician, reflecting local clinical practice. The laboratory also receives samples from patients from other hospitals around Vientiane; Friendship, Children’s and Setthathirat Hospitals. Written informed consent was obtained from patients or responsible guardians. Ethical clearance was granted by the Ethical Review Committee of the Former Faculty of Medical Sciences, National University of Laos and the Oxford University Tropical Ethics Research Committee. The confirmed aetiology was determined by the results of a panel of diagnostic tests which included tests for the direct detection of pathogens in CSF or blood, specific IgM in CSF, seroconversion, or a 4-fold rise in antibody titre between admission and follow-up serum samples (21). Pathogen detection was confirmed after critical analysis of test results to rule out possible contamination. Japanese encephalitis virus infection was confirmed, as recommended by the World Health Organisation, by detection of anti-JEV IgM by ELISA in CSF or seroconversion in paired serum samples. All anti-JEV IgM positive samples were subsequently confirmed by the gold standard virus neutralisation assay see cited reference (23). Power analysis was performed to estimate the sample size that would be required using different values. A schematic representation of the study methods is illustrated in Figure 1.

**Figure 1:**
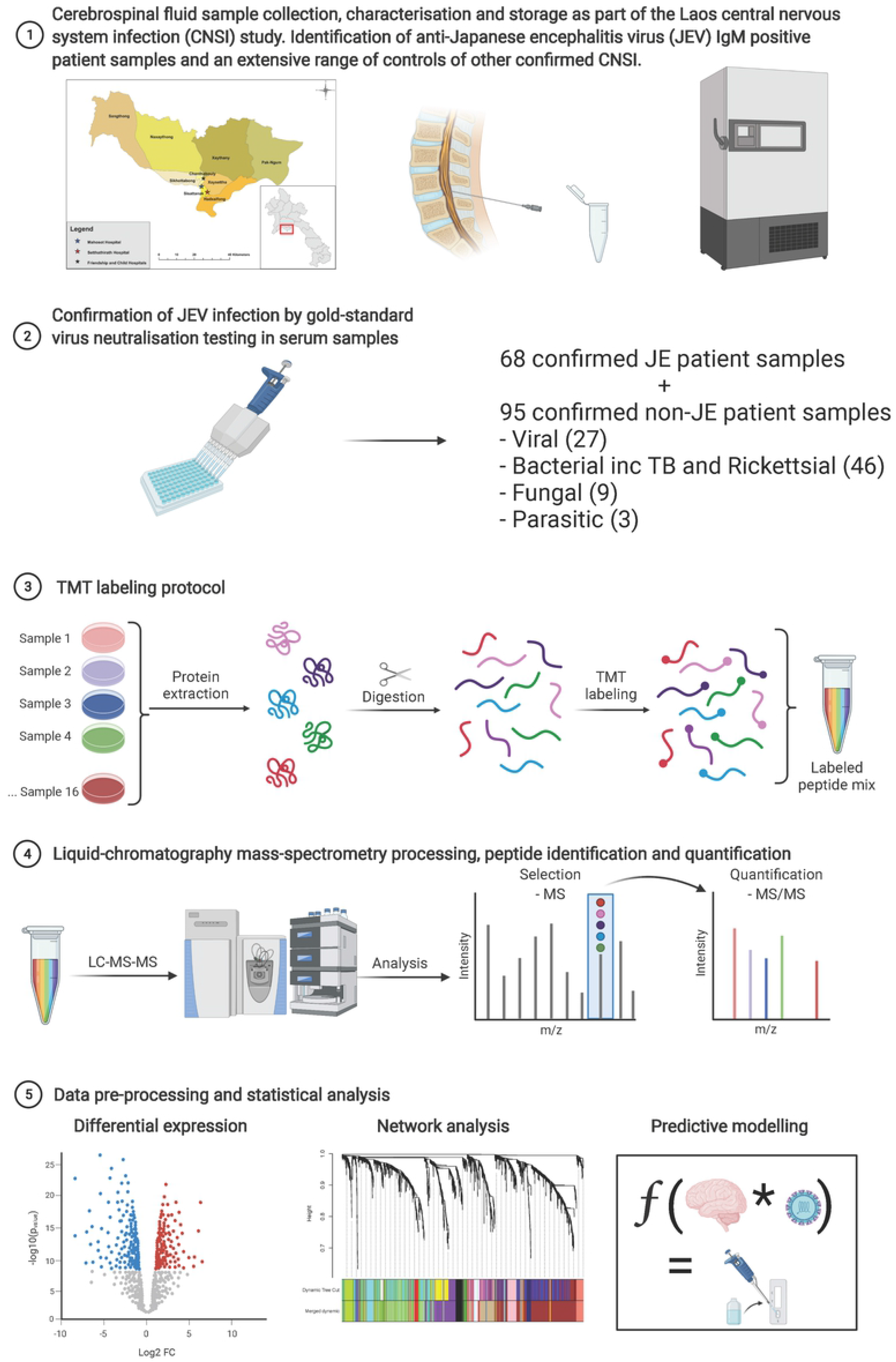
Schematic representation of the study methods

### LC-MS sample preparation

CSF samples were diluted 1:5 in 9 M urea and vortexed intermittently at room temperature for 30 minutes, to solubilise and denature proteins, inactivating any pathogens and rendering the sample acellular. Protein concentration was assessed with a Nanodrop assay ND-1000 spectrophotometer (Thermo Scientific) by measuring the absorbance at 280 nm, normalised by aliquoting different volumes of each sample dependent on the protein concentration, and then the total volume equalised with 7.5 M urea. An equal volume of 100 mM dithiothreitol (DTT) in 50 mM ammonium bicarbonate (AmBic) was added as a reducing agent, and the samples vortexed and incubated at 56°C for 45 min. An equal volume of 100 mM iodoacetamide (IAA) in 50 mM AmBic was added as an alkylating agent, vortexed and incubated at room temperature for 1 hr in the dark. 50 mM AmBic was added to each sample to reduce the urea concentration to below 1M. Digestion was performed with trypsin in a ratio of 1:20 m:m protein:trypsin (Promega, P/N V5072 for the pilot study; V5117 for the larger study); first 75% of the total amount of trypsin added and incubated at 37°C for 18 hours overnight and then the remaining 25% added and incubated at 37.5°C for 6 hours. The samples were frozen at - 20°C to quench the trypsin digestion reaction. A pooled aliquot of each sample was analysed by label-free LC-MS to verify protein digestion.

Reverse phase (RP) C18 solid phase extraction (SPE) was used to desalt the digested proteins, as per the manufacturers’ instructions (Waters P/N WAT023590 for the pilot study; Thermo Scientific P/N 60109-001 for the larger study). The total eluate was dried completely using a vacuum concentrator (Savant SpeedVac or Eppendorf concentrator) and for the samples to be labelled by Tandem Mass Tag (TMT), resuspended in 100 mM triethylammonium bicarbonate (TEAB). The samples were vortexed, centrifuged, sonicated for 3 min, and then this was repeated. The Pierce Quantitative Colorimetric Peptide Assay (Thermo Scientific, UK) was performed as per the manufacturer’s instructions. The samples were normalised for peptide concentration with TEAB to make up a final volume of 100 μL required for TMT labelling. TMT labelling was performed as per the manufacturer’s instructions, in two batches of TMT 11-plex (Thermo Scientific, P/N A37724) for the pilot study and ten batches of 16-plex (Thermo Scientific, P/N A44520) for the larger study. For the larger study, in order to examine technical variability and adjust for batch effects, each batch contained one reference pool and the batch 9 and 10 had two replicate samples. A pooled sample was analysed by LC-MS to verify labelling efficiency.

### Offline high pH reverse-phase fractionation

For the pilot study, offline high pH reverse-phase fractionation was performed using a Hypersil Gold column (Thermo Scientific, P/N 25002-202130). The mobile phase A was water adjusted with ammonium hydroxide to pH 10 and B was 10 mM ammonium bicarbonate in 80% Acetonitrile (ACN) adjusted with ammonium hydroxide to pH 10 and a flow rate of 300 µL/min. The samples were separated into 91 fractions with each fraction collected every 60 seconds from the start of the run and using the gradient shown in supplementary data (S1 Data). For the larger study, offline high pH reverse-phase fractionation was performed using an Xbridge BEH C18 column (Waters P/N 186006710). The mobile phase A was water adjusted to pH 10 with ammonium hydroxide and B was 90% ACN adjusted to pH 10 with ammonium hydroxide, at a flow rate of 200 µL/min. Fractions were collected every 60 seconds from the start of the run (100 fractions) and then concatenated into 44 fractions using the gradient shown in supplementary data (S1 Data). The samples analysed by DIA LC-MS/MS were not processed by offline fractionation.

### Liquid-chromatography mass-spectrometry

Online peptide desalting was performed with a Dionex Ultimate 3000 nano UHPLC (Thermo Scientific) using 100% of loading mobile phase A = 0.05% TFA in water at a flow rate 10 µL/min for 4.6 min. The online desalting column (trap column) used was a C18 column (Thermo Scientific P/N 160454). At 4.6 min the flow from the nano pump was diverted to the trap column in a backward flush direction. For online low-pH reverse-phase fractionation, the trapped peptides were eluted from the column over the gradient time specified in supplementary data (S1 Data). For the pilot study, Accucore C18 columns (Thermo Scientific P/N 16126-507569) were used with a nano source, at a flow rate of 250 µL/min. For the larger study, EASY-Spray PepMap C18 columns (Thermo Scientific P/N ES903) were used with an EASY-Spray source, and a flow rate of 300 µL/min. Mobile phase A was 0.1% FA and B was 0.1% FA in 80% ACN. MS was performed with a Q Exactive benchtop hybrid quadrupole-Orbitrap MS (Thermo Scientific), the settings are described in detail in supplementary data (S1 Data). For the CSF samples processed by DIA LC-MS/MS, samples were analysed using a Dionex Ultimate 3000 nano UPLC (Thermo Scientific) coupled to an Orbitrap Fusion Lumos mass spectrometer (Thermo Scientific). Briefly, peptides were trap on a PepMap C18 trap columns (Thermo) and separated on an EasySpray column (50cm, P/N ES803, Thermo) over a 60-minute linear gradient from 2 % buffer B to 35 % buffer B (A: 5 % DMSO, 0.1 % formic acid in water. B: 5 % DMSO, 0.1 % formic acid in acetonitrile) at a flow rate of 250 nL/min. The instrument was operated in data-independent mode as previously described (24).

### Data processing and statistical analysis

The sample size was estimated using a power calculation based on a t test and multiple testing correction, with data from the pilot study and the R package ‘FDRsampsize’ (25).

Protein identification, quantification, missing value imputation and batch correction: Thermo raw files were imported into Proteome Discoverer v2.5 (Thermo Scientific, UK) for peptide identification using the SEQUEST algorithm (26) searching against the SwissProt Homo sapiens and pathogen databases according to the included samples with precursor mass tolerance 10ppm and fragment mass tolerance 0.02 Da. Carbamidomethylation of cysteine, TMT at N-termini and lysine were set as fixed modifications, and oxidation of methionine was set as a variable modification. False discovery rate (FDR) estimation was performed using the Percolator algorithm (27). The criteria for protein identification included FDR < 1%, ≥ 2 peptides per protein, ≥ 1 unique peptides per protein, ≤ 2 missed cleavages and ≥ 6 and ≤ 144 peptide length (amino acids), coisolation threshold < 50%, average S/N threshold >10 and at least 2 channels with quantification data. Protein quantification was performed in R v 4.1.2 with the package MSstatsTMT (28). Proteins with >50% missing data were removed and the data was imputed with the package DreamAI (29). To incorporate peptide count per protein, jitter was added proportional to 1/median peptide count for each protein. The pilot and larger study data were merged, normalised with the package RobNorm (30) and then batch correction was performed with the function ComBat (31) in the package sva without modifiers as covariates (32). The protein list was filtered to remove contaminant proteins from the skin or red blood cells, see supplementary data S5_contaminants for the list of proteins removed.

Differential protein expression: Differential expression between the protein abundance in the JE vs. non-JE patient samples was performed using a t test and Benjamini-Hochberg correction for multiple testing.

Network analysis: Weighted correlation network analysis (WGCNA) was performed using the package WGCNA: constructing a signed weighted co-expression network with a soft power threshold of 12 to produce a power distribution, that is, scale-free topology; applying hierarchical clustering to detect modules of highly interconnected proteins with a minimum module size of five, deepSplit 4 and merge threshold 0.3; classifying intramodular hub proteins as the five proteins with the highest module membership for each module; and then correlating the modules with patient sample data (33).

Feature selection and predictive modelling: This was implemented with the Boruta algorithm (using the random forest classifier) using the package Boruta (34) and with Lasso (least absolute shrinkage and selection operator) regression using the package glmnet (16, 35). A final list of proteins based on the intersect between Boruta and Lasso were selected (36). Classification of JE vs. non-JE was performed with selected proteins using a several different machine learning models (random forest, support vector machine, logistic regression and naïve bayes with the package caret and caretEnsemble) (37). Models were trained using tenfold cross-validation repeated ten times evaluated on AUC-ROC. An analysis of feature importance was performed to identify proteins that best predicted the outcome (alive/ died) in JE patients, however due to the small sample size this was considered an exploratory analysis. Feature selection was performed with Boruta and Lasso, and then five-fold cross-validation performed on the entire dataset using different machine learning models. Protein involvement in biological, molecular and cellular processes was explored using gene ontology using the webserver STRING (38), the R package WebGestaltR 0.4.4 (39), and tissue expression correlated with the Human Protein Atlas (HPA) (40, 41).

Data independent acquisition (DIA) data processing: For robustness, final verification was performed on 10% of the samples independently processed via a separate mass spectrometry pipeline using label-free DIA LC-MS/MS. DIA data were analysed using DIA-NN software (v0.8) with the library-free approach as previously described (42), using the default settings as recommended. Briefly, for the library-free processing, a library was created from human UniProt SwissProt database (downloaded 24/2/21 containing 20,381 sequences) using deep learning. Trypsin was selected as the enzyme (1 missed cleavage), with carboamidomethylation of C as a fixed modification, oxidation of Methionine as a variable modification and N-term M excision. Identification and quantification of raw data were performed against the in-silico library applying 1% FDR at precursor level and match between runs (MBR). The DIA-NN ‘report.proteingroup’ matrix output was further analysed. Missing values were imputed with half the minimum value for each protein. These data was used as a test set in the predictive model for the diagnosis of JE. In view of the small numbers of JE patients included in the test set and missing outcome data for these patients, this was not used to test the predictive model for JE outcome.

## Results

### Patient data

Power analysis was performed to estimate the sample size that would be required to compare differential expression of proteins in JE vs non-JE using different values: with 1000-3000 biomarkers to be tested, 50-150 finally verified, effect size 0.8, power 90%, false discovery rate <5%, the total sample size with an equal number of JE cases and non-JE controls, was 122. Overall, including the pilot and larger study, 163 patients were included – 68 JE and 95 Non-JE, see Table 1, supplementary data S2 and S3.

**Table 1:** Summary of included patients’ demographics, clinical presentations and details of diagnosis.

JE patients were confirmed by the assays with the highest diagnostic confidence; detection of JEV RNA, or detection of JEV IgM in CSF or by seroconversion and confirmed by virus neutralisation tests (VNT). Non-JE patients included a range of different categories of infection that are common in the region. None of the patients had dual infections. Details of patient demographics, clinical presentations, laboratory investigations and outcome are reported in supplementary data S1 and S2.

### Protein profiling in CSF reveals differential expression in JE

5,070 proteins were identified, including 4,805 human proteins and 265 pathogen proteins, see supplementary data S4 for MSstatsTMT output for the pilot and larger studies. The pathogen proteins were bacterial or parasitic proteins. 2244 human proteins were identified in more than half of the samples included in both the pilot and larger studies. 68 proteins deemed to be contaminants were removed from the list, see supplementary data S5, resulting in a filtered list of 2176 proteins. 268 proteins showed differential expression (167 > 1.2 fold change, FC, and 101 <0.8 FC) based on the performance of a t test and Benjamini Hochberg multiple testing correction with p value <0.05, illustrated by the volcano plots in Figure 2.

**Figure 2:**
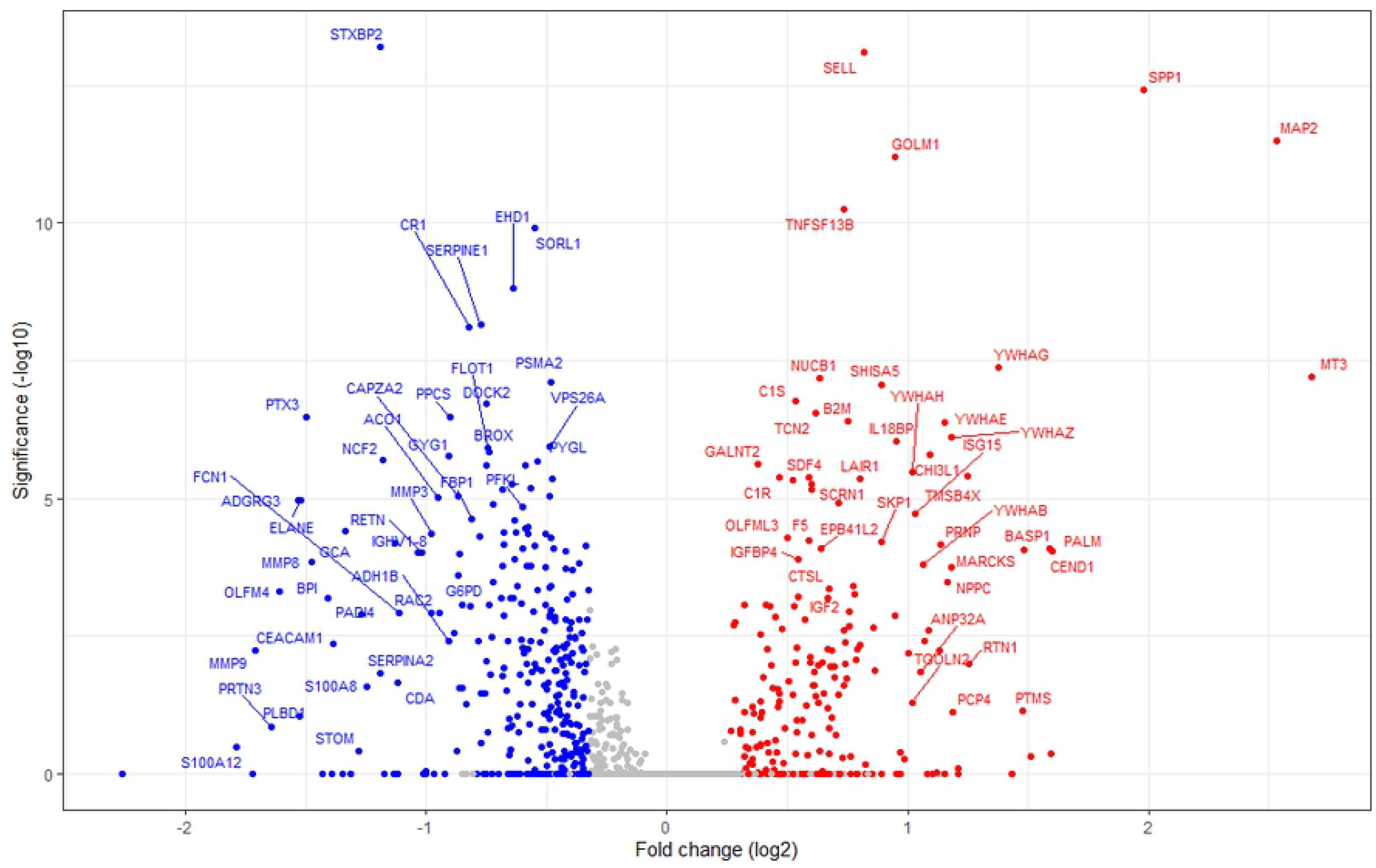
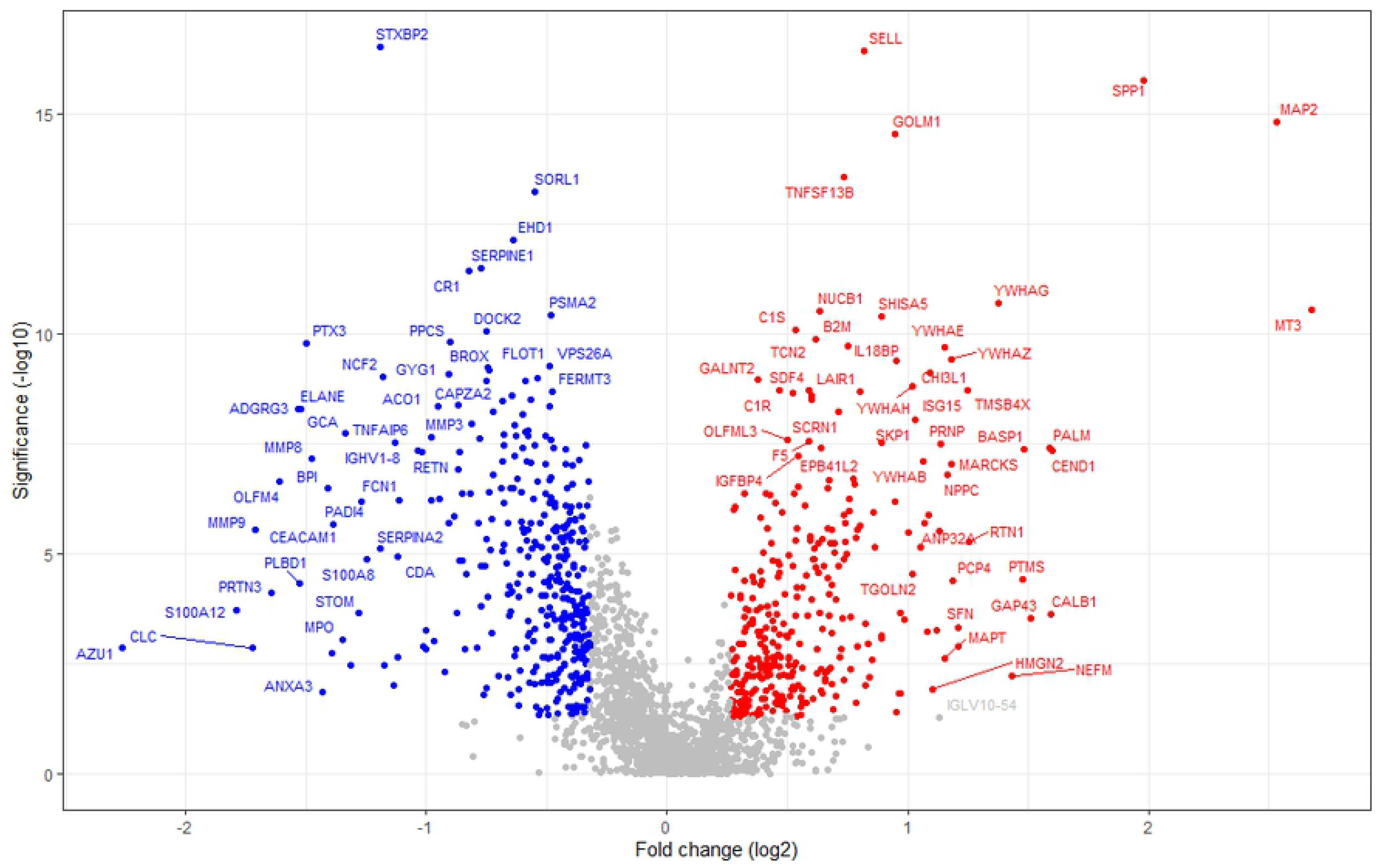
Volcano plots of the identified proteins illustrating the statistical significance (t test p values, a. uncorrected and b. corrected) against the magnitude of change (fold change) for Japanese encephalitis (JE) vs. Non-JE neurological infections.

### Molecular pathways associated with JE in CSF

2176 proteins from 163 patient samples were used to build a weighted gene expression network. A single outlier was identified, see supplementary data S7, and removed. Further analysis revealed that this sample had higher overall protein abundances, in spite of peptide normalisation prior to TMT labelling and downstream normalisation in MSstatsTMT and RobNorm during data processing. 44 modules were identified, and then closely related modules merged into 20 modules, see the tree diagram illustrating the cluster dendrogram in Figure 3 and the modules in Figures 4. Module-trait relationships are shown in Figure 5; suggesting that 15 modules were associated with JE (p value < 0.05), 9 upregulated (red) and 6 downregulated (green). 10 of the modules included proteins in the top five intramodular proteins, that is proteins with the highest modular membership, with significant differences in abundance between the JE and non-JE group.

**Figure 3:**
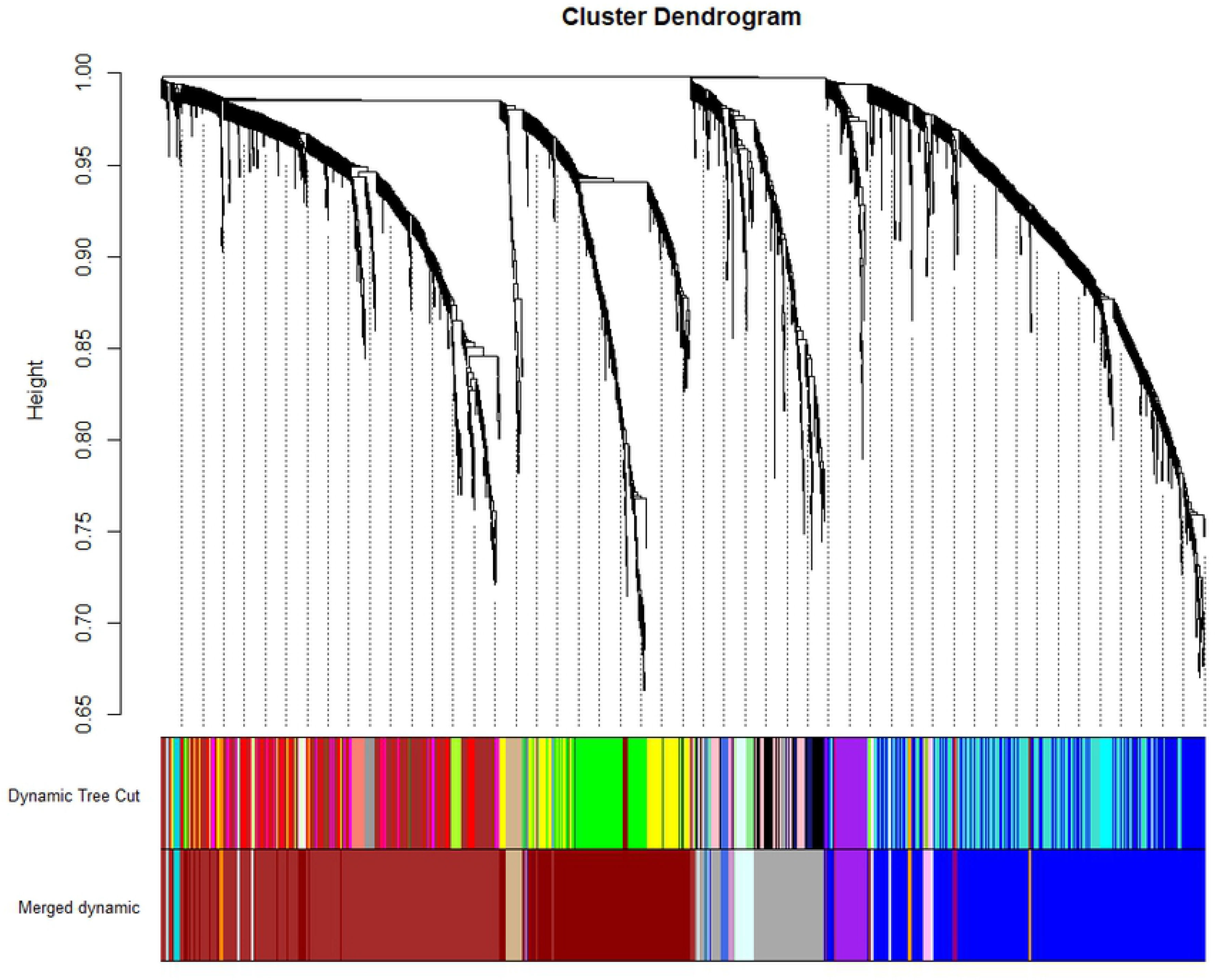
Weighted correlation network analysis cluster dendrogram

**Figure 4:**
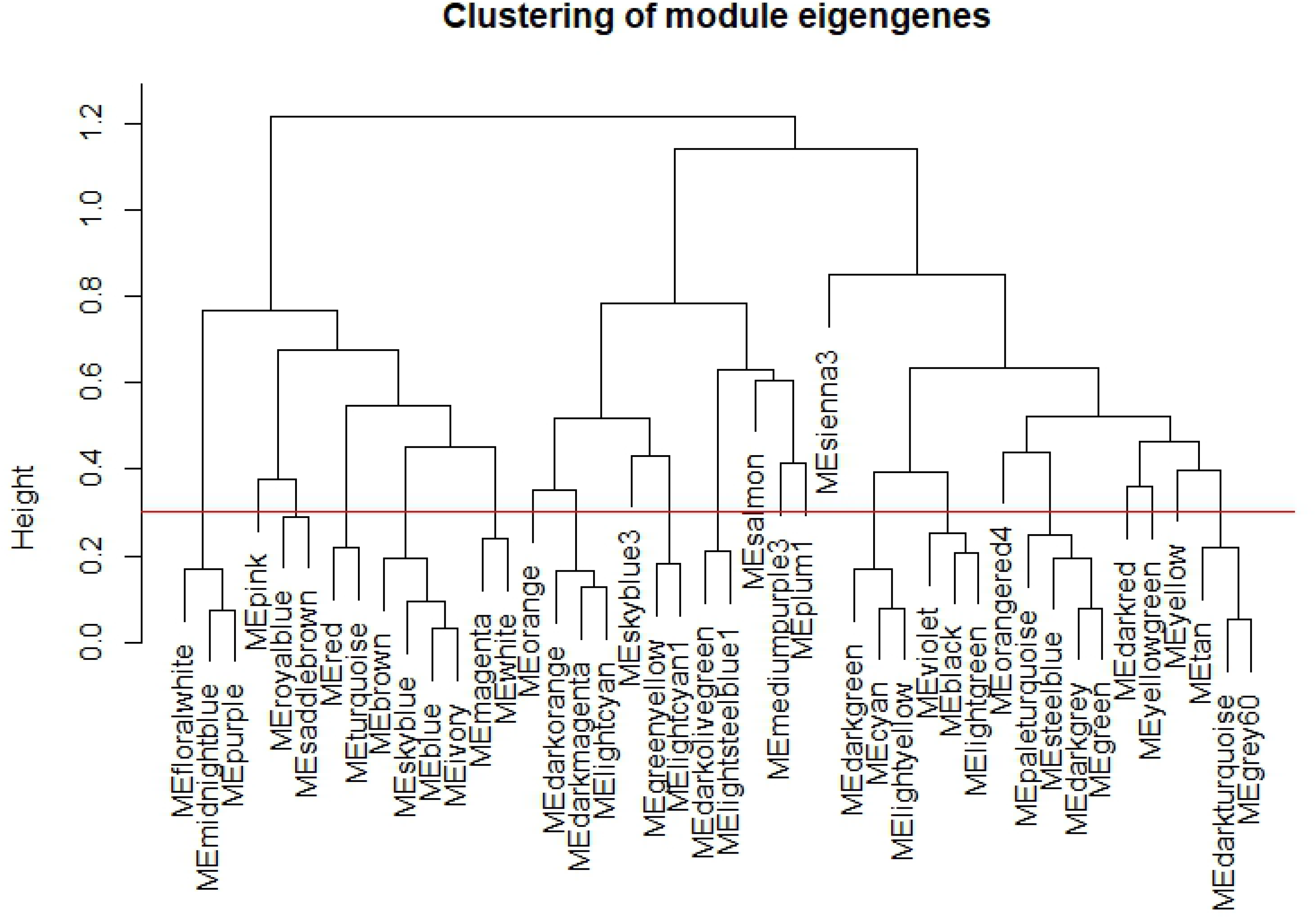
Weighted correlation network analysis clustering of module eigengenes. The red line in the figure indicates the threshold for merging modules together, here the threshold was 0.3.

**Figure 5:**
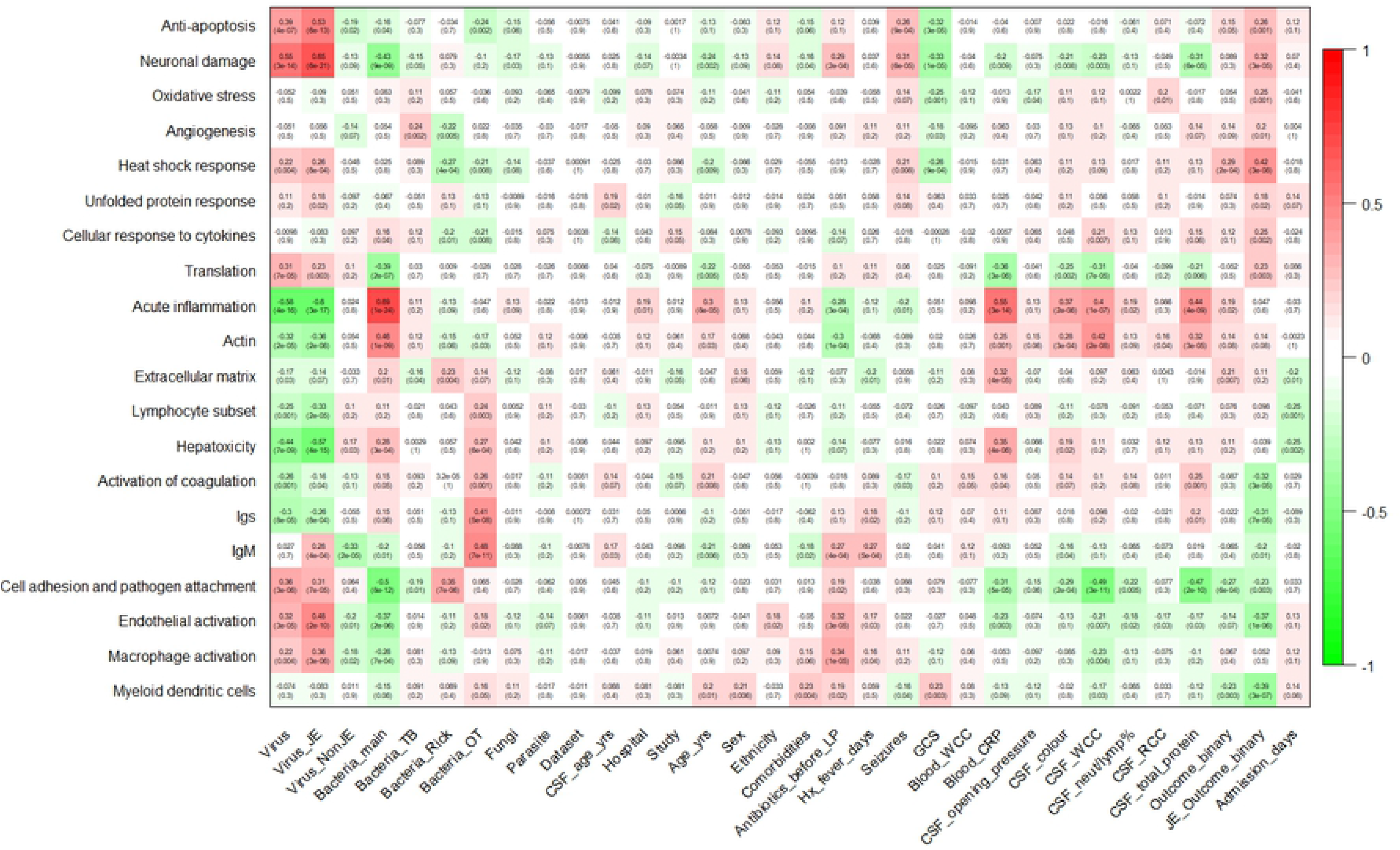
Weighted correlation network analysis module-trait relationships. darkred=anti-apoptosis, red=neuronal damage, sienna3=oxidative stress, orangered4=angiogenesis, yellowgreen=heat shock response, yellow=unfolded protein response, darkgreen=cellular response to cytokine, floralwhite=translation, darkolivegreen=acute inflammation, paleturquoise=actin, salmon=extracellular matrix, mediumpurple3=lymphocyte subset, plum1=hepatotoxicity, darkorange=activation of coagulation, greenyellow=Igs, skyblue3=IgM, brown=cell adhesion and pathogen attachment, magenta=endothelial activation, pink=macrophages, royalblue=myeloid dendritic cells.

### A diagnostic protein signature of JE in CSF

Feature selection: In total, 86 proteins were identified by at least one of the feature selection procedures as important in classifying JE vs non-JE; 68 proteins identified with the Boruta algorithm and 28 with Lasso, see supplementary data S10. The proteins were associated with 11 different WGCNA modules, all of which had been identified as associated with JE through WGCNA. 48 were upregulated and 38 downregulated in comparison to other neurological infections. Functional enrichment analysis in STRING demonstrated interactions between the proteins, Figure 6. Gene ontology analysis highlighted overexpression of proteins related to apoptosis and downregulation of proteins related to neutrophil degranulation, supplementary data S11. 22 proteins were secreted proteins: Immunoglobulin lambda variable 3-9 (IGLV3-9), Immunoglobulin heavy variable 3-74 (IGHV3-74), Golgi membrane protein 1 (GOLM1), Cathepsin L (CTSL), CEA cell adhesion molecule 8 (CEACAM8), Phospholipase B domain containing 1 (PLBD1), Cerebellin 1 precursor (CBLN1), Secreted phosphoprotein 1 (SPP1), Natriuretic peptide C (NPPC), Microtubule associated protein tau (MAPT), Chitinase 3 like 1 (CHI3L1), ISG15 ubiquitin like modifier (ISG15), Interleukin 18 binding protein (IL18BP), Beta-2-microglobulin (B2M), TNF superfamily member 13b (TNFSF13B), Bactericidal permeability increasing protein (BPI), Pentraxin 3 (PTX3), Matrix metallopeptidase 9 (MMP9), S100 calcium binding protein A12 (S100A12), Azurocidin 1 (AZU1), Olfactomedin 4 (OLFM4) and Matrix metallopeptidase 8 (MMP8). 15 proteins were associated with increased expression in the brain: Brain abundant membrane attached signal protein 1 (BASP1), Aldolase, fructose-bisphosphate C (ALDOC), CBLN1, Metallothionein 3 (MTX3), MAP2, Tyrosine 3-monooxygenase/tryptophan 5-monooxygenase activation protein gamma (YWHAG), Tyrosine 3-monooxygenase/tryptophan 5-monooxygenase activation protein eta (YWHAH), MARCKS like 1 (MARCKSL1), Secernin 1 (SCRN1), SPP1, Microtubule associated protein tau (MAPT), CHI3L1, Paralemmin (PALM), Reticulon 1 (RTN1), Purkinje cell protein 4 (PCP4), Cytidine/uridine monophosphate kinase 2 (CMPK2), NPPC, Glial fibrillary acidic protein (GFAP), Cell cycle exit and neuronal differentiation 1 (CEND1). Thus, three proteins were secreted and showed an increased expression in the brain: SPP1, MAPT, CHIL3, NPPC and CBLN1.

**Figure 6:**
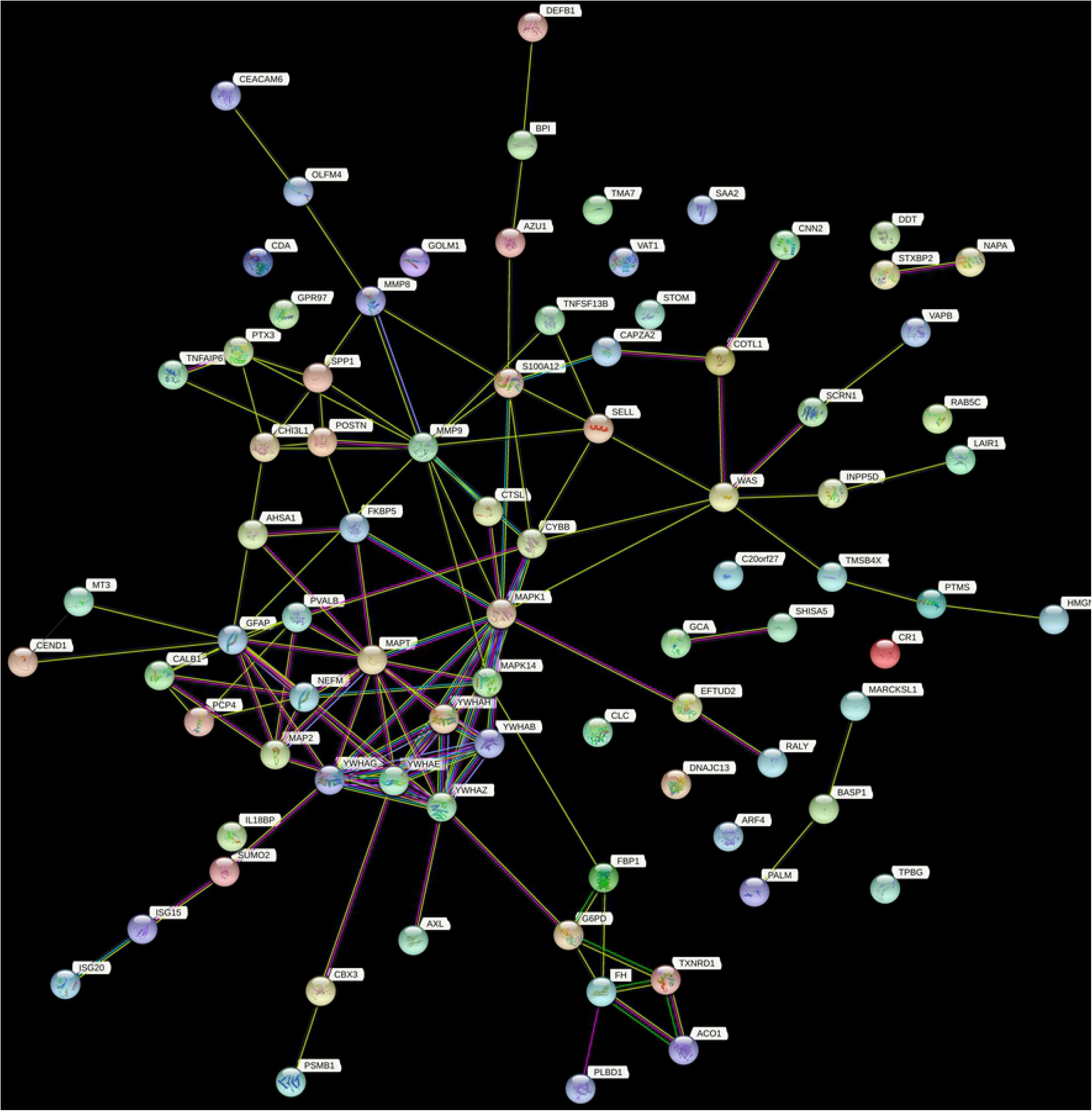
STRING functional protein association network https://version-11-5.string-db.org/cgi/network?taskId=bMZcY3ZdJua4&sessionId=bpgLGeFd4RM1

JEV has a predilection for the thalamus and substantia nigra of the basal ganglia (23). One of the proteins were ‘group enriched’ in the thalamus, MMP9, from the HPA database. Four proteins were associated with the GO term ‘substantia nigra development’, associated with BASP1, Glucose-6-phosphate dehydrogenase (G6PD), YWHAH and 14-3-3 protein epsilon (14-3-3epsilon). The HPA database includes mRNA expression data from 13 brain regions, including the basal ganglia and thalamus; substantia nigra expression on its own is not reported (https://www.proteinatlas.org/humanproteome/brain).

#### Predictive modelling

Feature selection identified a final set of 10 proteins which together exhibited high predictive performance (Figure 7). When examined using the ensemble model, using ten-fold cross validation, JE classification demonstrated an AUC-ROC of 99.5 (99.2-99.9), in addition to high sensitivity and specificity – metrics in Table 2 and ROC in supplementary data S12.

**Figure 7:**
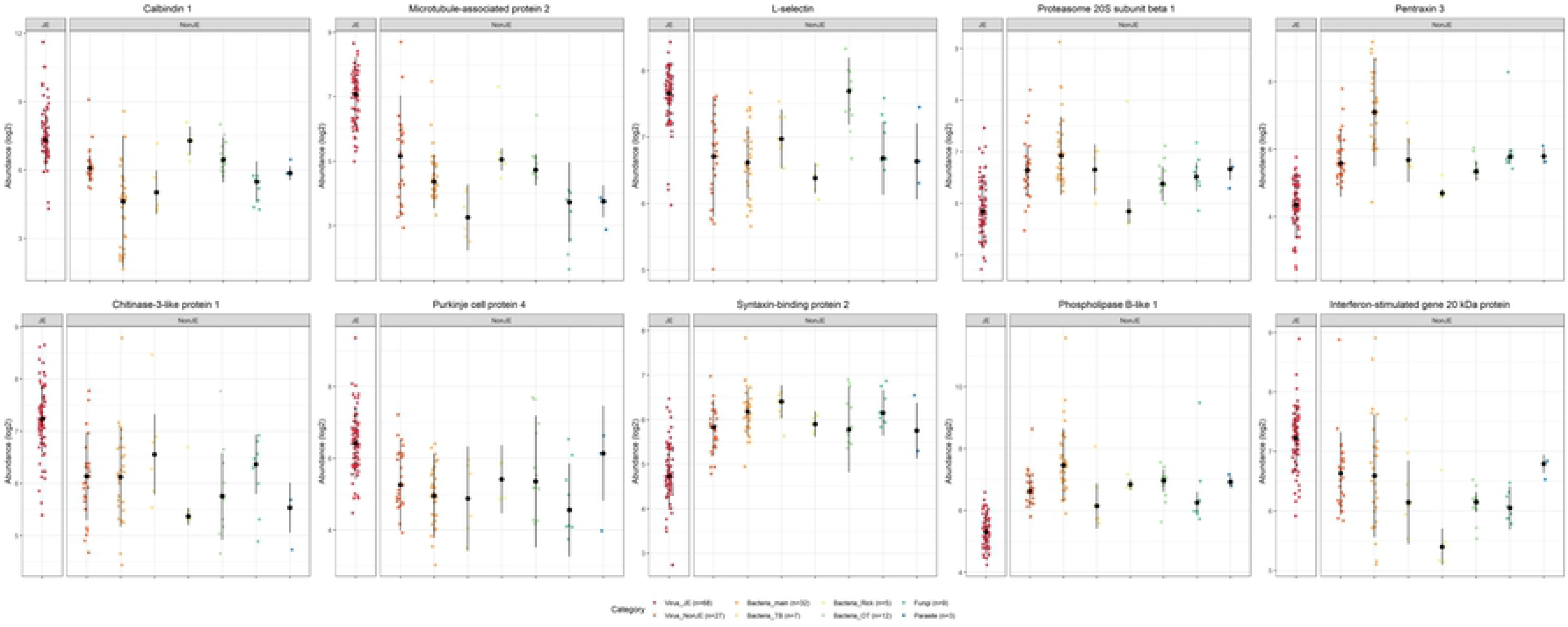
Differential expression across samples in ten proteins as a diagnostic signature of Japanese encephalitis virus infection

**Table 2:** Predictive modelling scores with 95% confidence intervals

| Classification task | Data | AUC-ROC | Accuracy | Sensitivity | Specificity | Positive predictive value | Negative predictive value |
| --- | --- | --- | --- | --- | --- | --- | --- |
| JE diagnosis (JE vs. non-JE) | Training set<sup>1</sup> (n=163) | 99.5 (99.2-99.9) | 99.4 (98.8-99.8) | 99.5 (98.5-99.9) | 99.3 (98.3-99.8) | 99.3 (98.3-99.8) | 99.5 (98.5-99.9) |
|  | Test set<sup>2</sup> (n=16) | 91.0% (79.0-100) | 87.5% (61.7-98.5) | 100% (47.8-100) | 81.8% (48.2-97.7) | 71.4% (29.0-96.3) | 100% (66.4-100) |
| JE outcome (dead vs. alive) | Training set<sup>3</sup> (n=42) | 88.5 (84.7-92.2) | 86.3 (83.5-88.8) | 42.0 (32.2-52.3) | 93.7 (91.4, 95.5) | 52.5 (41.0-63.8) | 90.6 (88.1-92.8) |

1. The training set included patient samples (68 JE and 95 non-JE confirmed neurological infections) processed by TMT LC-MS/MS. 2. The test set included 10% of the patient samples from the TMT LC-MS/MS analysis processed separately by label-free DIA LC-MS/MS. 3. The training set included all the JE patients included in the TMT LC-MS/MS analysis for which outcome data was available.

Data acquired by DIA LC-MS/MS of 16 (10%) of the samples was used to verify the ten-protein JE diagnostic predictive model. The test metrics are reported in Table 2.

### Predictors of JE outcome

Feature selection: Subgroup analysis was performed using 42 JE samples for which outcome data at hospital discharge (died vs. alive) were available. Seven proteins were identified as important in predicting outcome using the Boruta algorithm and 2 proteins using Lasso, such that 2 proteins were identified by both Boruta and Lasso, see supplementary data S13. In view of the small sample size, the data were not split into a training and test set. These proteins were used to train different models with five-fold CV repeated ten times evaluated on ROC, and then combined in an ensemble model with cross-validation scores reported in Table 2, see the list of proteins in supplementary data S13 and ROC in S14. There were five JE patients in the DIA LC-MS analysis of which 3 had outcome data, and this was considered too small to report test metrics.

## Discussion

We performed deep untargeted analysis of well-characterised patient CSF samples from a large number of different confirmed neurological infections. To our knowledge, the highest number of proteins in CSF identified to date has been 3,174 (43); thus this research represents a notable improvement in terms of the numbers of proteins identified and this serves as a marker of the depth of analysis and prospects for biomarker identification (44). Offline fractionation into 90 fractions in the pilot study, and 100 fractions concatenated into 44 in the larger study, with two-hour online LC gradients and multiplexing with TMT-16plex contributed to the depth of analysis. Furthermore, the diverse range of neurological infections also augmented the variety of proteins identified.

WGCNA analysis identified 20 clusters of highly correlated proteins, and provided insight into the proteins and how they associate with disease mechanisms. The modules were allocated a descriptor, according to functional enrichment analysis of the proteins. For example, one module was associated with IgM (proteins in the module included Immunoglobulin heavy constant mu and Immunoglobulin J chain) and correlated with JE and *Orientia tsutsugamushi* (OT), as well as the duration of illness. Other important modules associated with upregulation in JE included neuronal damage, anti-apoptosis, heat shock response, unfolded protein response, cell adhesion, macrophage and dendritic cell activation. In contrast, in comparison to other non-JE neurological infections, there was an association with downregulated acute inflammatory response, hepatotoxicity, activation of coagulation, extracellular matrix and actin regulation.

Predictive modelling using the 10 protein ensemble model enabled classification of JE and non-JE samples with a CV accuracy of 99.4 (95% CI 98.8-99.8) across all the samples using the TMT labelled DDA data, and 87.5% (95% CI 61.7-98.5) in verification with 16 (10%) of the samples by DIA. DIA is a label-free method of analysis, with ongoing improvements in depth and throughput; in this case providing a complementary method to verify the TMT data rather than performing traditional targeted LC-MS/MS proteomics such as parallel reaction monitoring (PRM). Three proteins selected as the best disease classifiers were not “significant” i.e. p value < 0.05 with t-test and adjustment for multiple testing, highlighting the limitations of univariate analysis in biomarker identification (45). Biomarker discovery is a lengthy process, akin to the pharmaceutical pipeline (13). The work demonstrates important CSF proteins in classifying JE vs. non-JE. However, there is no doubt that the protein signature needs to be validated with orthogonal antibody-based methods in additional patient groups. It will also be useful to compare this with protein profiling in other body fluids. This will inform the use of a smaller subset of proteins in an ELISA or rapid diagnostic test (RDT) to be tested alongside the existing anti-JEV IgM assay.

To date, to our knowledge, two studies have utilized unbiased techniques to examine the CSF proteome in human patients with confirmed JEV infection; while they demonstrates the feasibility of the methods, the patients were not confirmed by seroneutralisation and included relatively small numbers of patients (10 and 26 JE patients) (46, 47). There have been a handful of studies utilizing ELISA methods to target specific proteins, however these rarely used power calculations in their experimental design, nor did they include adequate controls (48–53). Analysis of the transcriptome and proteome in animal models (54–58) and cell culture (48, 54, 59–64) have been performed, however the comparability to human CSF and comparison with other neurological infections is limited. Furthermore, mRNA expression does not directly correlate with that of the corresponding protein (65).

As expected, while we included JEV proteins in the search database, we did not identify any JEV pathogen proteins. This is compatible with previous publications; non-structural protein 1 is the major secreted protein during flavivirus infections, harnessed widely as a diagnostic biomarker for Dengue virus infection, but not a useful diagnostic biomarker for JE (66). The data provide useful interrogation of the host response to JEV infection. The identified proteins fit well into the existing literature on the host response in JEV and other closely-associated flavivirus infections, most importantly West Nile virus infection (67, 68). MAPT and MAP2 are both closely associated microtubule stabilising proteins specific to neuronal cells (69). Both proteins were identified in this study as being biomarkers of JE in CSF, and the high levels in comparison to other neurological infections is striking. The association of the former has previously been demonstrated by ELISA, in one of the only studies of this type (70). The role of actin, microtubule and intermediate filament cytoskeletal re-organisation in flavivirus infection has been described (71) and upregulation of MAPT and MAP2 may represent neuronal damage following transneural spread of JEV. Other proteins that were associated with JE in this study, all within the red WGCNA module, that may reflect neuronal damage include Paralemmin, Calbindin 1, MAP2, Parvalbumin, Secernin 1 and Cell cycle exit and neuronal differentiation. The upregulation of ISG15 and ISG20 fit in with the known upregulation of a host of ISGs as part of the innate immune response to a viral infection (72, 73). Additional functional enrichments reflecting different WGCNA modules have previously been described anti-apoptosis (74), heat shock response (75, 76), unfolded protein response (77), translation (78), IgM (79), cell adhesion and pathogen attachment (80), endothelial activation (81) and macrophage activation (82, 83). In comparison to other neurological infections, there was a downregulation in acute phase response proteins and neutrophil enriched proteins, as has been seen by other studies (84–87). In these, however, the sample size for the analysis of proteins predictive of outcome was less substantial and not supported by an a priori power calculation.

Incomplete coverage and missing data between LC-MS runs is an ongoing issue in the field (29). It is notable that comparing with other similar studies in the literature, the important proteins may not be exactly the same but are closely related. These issues are now being improved by DIA methods. Further limitations are that we did not include CSF from healthy people in Laos on ethical grounds, or from cohorts from elsewhere on the basis that samples that have undergone different storage conditions may not be comparable. The latter is also the reason that there are no samples from neurological flaviviruses occurring in other geographical areas, such as West Nile virus (WNV) and Zika virus (ZIKV). Furthermore, for the purposes of the objective of finding a diagnostic protein signature of JE, the utmost importance was comparing JE with controls of a wide range of other neurological infections. The analysis of proteins predictive of different categories of infectious aetiologies was not sufficiently powered, and has not been reported. It is important to keep in mind that the comparison is between different neurological infections in the analysis of proteins that are up and down-regulated.

An RDT to detect JE in less accessible areas is urgently needed. This study demonstrates the feasibility of an unbiased LC-MS approach in the identification of novel protein biomarkers of neurological infections. Additional data using antibody-based methods will allow the 10-protein signature to be refined. This will require purchasing or development of ELISA assays and comparing the specific protein abundance in JE and non-JE patients. These data will need to be validated in a larger group of patients, in different locations and in field settings. Ultimately, this will enable the selection of 2-3 proteins for the development of an RDT.

## Author contributions

TB, BG, NZ, ADP and PNN conceived the study. The Laos CNS study was completed by ADP, PNN, XDL, MV, MM, SP, AC, OS, OP and the SEAe collaborators. JEV seroneutralisation was performed in Marseille by TB, NA, BP and supervised by ADP and XDL. TB, BG and NZ developed the methodology for the TMT LC-MS/MS analysis, and TB performed the laboratory work with input from AK, BG and DO. IV, RF and BK developed the methodology for the DIA LC-MS/MS analysis, and TB and IV performed the laboratory work. TB performed the data analysis with input from AM. TB wrote the manuscript; all the authors edited successive drafts and approved the final version.

## Acknowledgements

We are very grateful to the patients and to Bounthaphany Bounxouei, the former Director of Mahosot Hospital, the late Rattanaphone Phetsouvanh, Director of the Microbiology Laboratory, and the staff of the wards and Microbiology Laboratory of Mahosot Hospital. We also thank Bounnak Saysanasongkham, the former Director of the Department of Healthcare and Rehabilitation, Ministry of Health, and Bounkong Syhavong, Minister of Health, Lao PDR for their very kind help and support. We thank the stakeholders of the SEAe project, members of the Unité des Virus Émergents (Christine Isnard and Camille Placidi) and the CNR des Arbovirus (Patrick Gravier, Gilda Grard, Isabelle Leparc-Goffart and Mathilde Galla). Finally, we acknowledge useful discussions on the data analysis with Andrew R. Jones at the University of Liverpool, and Damien Ming, Mauricio Barahona and Robert Peach at Imperial College London.

## South-East Asia Encephalitis (SEAe) Study (SEAe) Collaborators

We are grateful to all the SEAe study researchers, including Philippe Buchy, Em Bunnakea, Julien Cappelle, Mey Channa, Veronique Chevalier, Yoann Crabol, Philippe Dussart, Marc Eloit, Christopher Gorma, Magali Herrant, Nguyen Hien, Chaw Su Hlaing, Jérôme Honnorat, Tran Thi Mai Hung, Tran Thi Thu Huong, Latt Latt Kyaw, Nguyen Van Lam, Denis Laurent, Marc Lecuit, Kyaw Linn, Olivier Lortholary, Aye Mya Min Aye, Philippe Perot, Sommanikhone Phangmanixay, Khounthavy Phongsavath, Phan Huu Phuc, Anne-Laurie Pinto, Patrice Piola, Jean-David Pommier, Bruno Rosset, Ky Santy, Heng Sothy, Arnaud Tarantola, Nguyen Thi Thu Thuy, Htay Htay Tin, Ommar Swe Tin, Pham Nhat An, Dang Duc Anh, Pascal Bonnet, Kimrong Bun, Danoy Chommanam, Viengmon Davong, Patrice Debré, Jean-François Delfraissy, Christian Devaux, Anousone Douangnouvong, Veasna Duong, Benoit Durand, Chanreaksmey Eng, Catherine Ferrant, Didier Fontenille, Lukas Hafner, Le Thanh Hai, Do Thu Huong, Marc Jouan, May July, Magali Lago, Jean-Paul Moatti, Bernadette Murgue, Khin Yi Oo, MengHeng Oum, Khansoudaphone Phakhounthong, Anh Tuan Pham, Do Quyen, Malee Seephonelee, Maud Seguy, Bountoy Sibounheunang, Kanarith Sim, Luong Minh Tan, Cho Thair, Win Thein, Phung Bich Thuy, Hervé Tissot-Dupont and Malavanh Vongsouvath.

## Funding

TB was supported by the University of Oxford and the Medical Research Council (grant MR/N013468/1). The work was also supported by the Oxford Glycobiology endowment, the Institute of Research for Development, Aix-Marseille University and the European Union’s Horizon 2020 research, Fondation Total, Institut Pasteur, International Network Institut Pasteur, Fondation Merieux, Aviesan Sud, Institut national de la santé et de la recherche médicale (Inserm), and innovation programme EVAg (grant agreement 653316). This research was funded in part by the Wellcome Trust. For the purpose of Open Access, the author has applied a CC BY public copyright licence to any Author Accepted Manuscript version arising from this submission. AM was supported by a scholarship from the Medical Research Foundation National PhD Training Programme in Antimicrobial Resistance Research (MRF-145-0004-TPG-AVISO). DPO, IV, RF and BK are supported by the CAMS China Oxford Institute. LT is a Wellcome clinical career development fellow, supported by grant number 205228/Z/16/Z, and the NIHR Health Protection Research Unit in emerging and zoonotic infections (NIHR200907) at University of Liverpool in partnership with Public Health England (PHE), in collaboration with Liverpool School of Tropical Medicine and the University of Oxford.

## Competing interests

None declared.

## Ethical approval

Ethical clearance for the Laos CNS study was granted by the Ethical Review Committee of the former Faculty of Medical Sciences, National University of Laos (now University of Health Sciences) and the Oxford University Tropical Ethics Research Committee, Oxford, UK.

## Data availability

The mass spectrometry proteomics data sets were submitted to the PRIDE public data repository. All other data underlying this article are available in the article and in its online supplementary material.

## Notes

### Competing Interest Statement

The authors have declared no competing interest.

